# Infauna selectively enhance DNA virus diversity and activity in marine sediments

**DOI:** 10.64898/2026.03.17.711404

**Authors:** Alexis Fonseca, Mathias Middelboe, Karin Holmfeldt, Emma Bell, Christoph Humborg, Alf Norkko, Francisco J.A. Nascimento

## Abstract

Viruses regulate microbial mortality and biogeochemical cycling in marine sediments; however, the ecological drivers of sediment viral communities remain unclear. Infauna, including sediment-dwelling meiofauna and macrofauna, are major ecosystem engineers that reshape sediment structures and microbial processes, but their influence on viruses is unknown. We combined infaunal gradient incubations with metagenomic and metatranscriptomic analyses to assess viral DNA and RNA responses. DNA viruses showed increased abundance (3-fold), diversity, richness, and transcriptional activity under higher infauna abundance conditions, whereas RNA viruses remained unaffected, revealing striking selectivity. This selectivity reflects an infauna-dependent component mediated by bacterial activity that cannot be explained by host abundance alone. Infection profiling revealed increased transcription of viral replication and structural genes, and lytic viruses under high infauna conditions. These findings establish infauna as a previously overlooked regulator of DNA virus dynamics, integrating viral ecology into faunal-microbial frameworks in benthic ecosystems and suggesting potential influences on geochemical cycles.

**Teaser:** Infauna selectively shape viral communities in marine sediments, revealing an overlooked effect on DNA viruses.

## Introduction

Viruses are the most abundant biological entities in aquatic environments (Suttle, 2007; Gregory et al., 2019). Marine waters contain, on average, ∼10^7^ viral particles per milliliter, whereas viral densities in coastal sediments range from 10^5^ to 10^9^ particles per cm^3^ (Wommack et al., 2000; Engelhardt et al., 2014). In marine sediments, viral production is tightly coupled to bacterial metabolic activity (Middelboe and Glud, 2006). Through host cell lysis, reprogramming of host metabolism, and expression of auxiliary metabolic genes (AMGs), viruses play a central role in microbial mortality, organic matter turnover, and sediment biogeochemical cycling (Chevallereau et al., 2022; Luo et al., 2022; Middelboe et al., 2025).

Classical frameworks in viral ecology emphasize bottom-up controls, whereby viral abundance and production are primarily governed by microbial host availability and productivity (Weinbauer, 2004; Danovaro et al., 2008). Consequently, bacterial abundance, activity, and diversity are major determinants of viral community structure in sediments (Middelboe et al., 2006). These microbial properties are shaped by environmental drivers, such as the availability of organic matter, oxygen conditions, and temperature, which collectively regulate benthic biogeochemical processes (Sanders and Kalff, 1993; Hicks et al., 2018). In addition to these physicochemical controls, bioturbation by infauna represents a powerful driver of the microbial diversity and structure in sediments. Infauna, comprising meiofauna (0.04-1 mm) and macrofauna (>1 mm), modify sediment structure through burrowing, feeding, and ventilation, thereby redistributing particles, solutes, and microorganisms (Aller, 1994; Kristensen et al., 2012; Bonaglia et al., 2013). Bioturbation by infauna is now recognized as a universal process in aquatic and terrestrial sediments that enhances microscale heterogeneity and regulates key biogeochemical cycles, including carbon, nitrogen, and methane (Meysman et al., 2006; Laverock et al., 2014; Broman et al., 2024).

Despite the central role of infauna in shaping sediment microbial communities, their influence on viral community structure and activity remains largely unexplored. This gap is particularly relevant given that viral communities comprise biologically distinct groups with fundamentally different genome types and host associations. DNA viruses in sediments are predominantly bacteriophages (Yu et al., 2024), which respond directly to bacterial abundance, while RNA viruses are mainly associated with eukaryotic hosts (Zhang et al., 2024). Given their distinct host associations, we therefore hypothesised that DNA and RNA viruses would respond differently to infaunal gradients in abundance, size and bioturbation intensity, where DNA viruses are stronger affected. Here, we employed infauna gradient experiments established by Broman et al. (2024), combined with metagenomic and metatranscriptomic approaches, to investigate how variations in infaunal abundance influence sediment viral communities in coastal sediments. By investigating DNA and RNA virus diversity and community structure across infaunal abundance and size gradients, we revealed that sedimentary viral communities respond selectively to such gradients. These findings position infauna as an overlooked ecological control of sediment viral ecosystems and underscore the integration of viruses into faunal-microbial interactions in aquatic systems.

## Results

### DNA and RNA virus abundance and diversity

#### DNA viruses

A total of 141 DNA virus genomes (Supplementary Data 1) were obtained and classified into 21 families (Table S1). In terms of relative abundance (adjusted coverage > 4x), the dominant family was *Peduoviridae* (59.6%), a potentially lysogenic dsDNA phage within *Caudoviricetes* (Fig. 1A), *Adenoviridae*-like (7.9%), dsDNA viruses with homologs across metazoan hosts, and *Inoviridae*-like (3.1%), which are positive-sense ssDNA phages, respectively. Of the remaining, 23.4% were labelled as unclassified. DNA virus abundance was significantly higher under high meiofauna (HM) and high meiofauna + macrofauna (HMM) conditions than under low meiofauna abundance (LM) (Fig. 1C). Similarly, the indices Shannon, Simpson diversity, and richness were significantly higher (*p* < 0.05) in HM and HMM than LM condition (Fig. 1D) (Table S2).

**Fig. 1.**
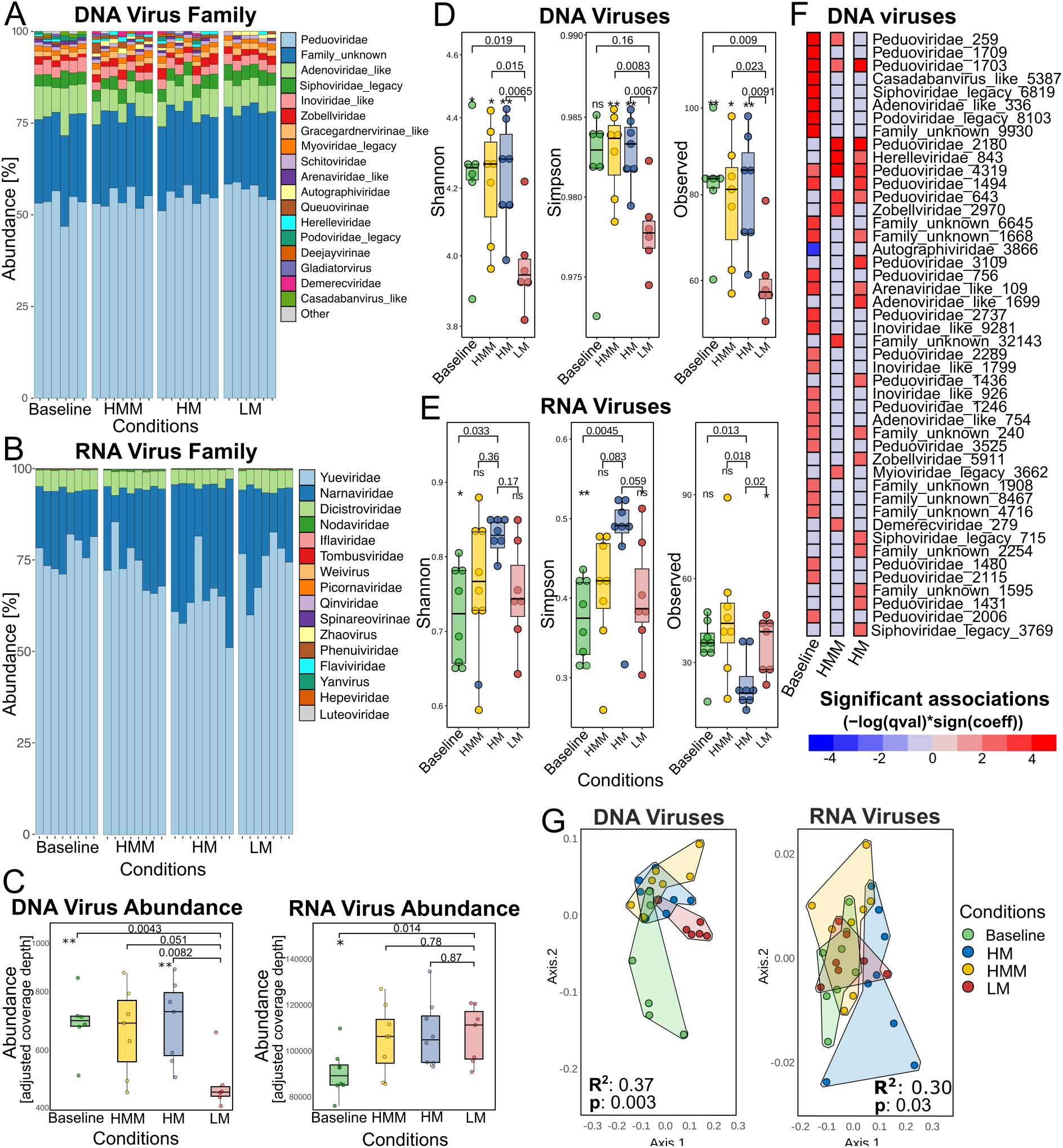
Taxonomy, abundance, and diversity of DNA and RNA viruses across infauna treatments. (**A**) Relative abundance of DNA virus families (n = 21). Abundance was based on the adjusted coverage depth. Taxa labelled as “legacy” are deprecated from the International Committee on Taxonomy of Viruses (ICTV). (**B**) Relative abundance of RNA virus families (n = 12). (**C**) Total abundance (cumulative adjusted coverage) of DNA and RNA viruses across the infauna treatments (Wilcoxon test, *p* < 0.05). (**D**) Diversity (Shannon and Simpson) and richness (observed) indices of DNA viruses (Wilcoxon test, *p* < 0.05). (**E**) Diversity and richness indices of the RNA viruses. (**F**) Differential abundance of DNA viruses (MaAsLin2, *p* < 0.05, *q* < 0.09) using LM as a reference. (**G**) Principal coordinates analysis (PCoA) of DNA and RNA viral communities based on relative abundance and Bray-Curtis dissimilarity (PERMANOVA Adonis, *p* < 0.05).

PCoA based on relative abundance and dissimilarity Bray-Curtis index showed that DNA viruses clustered according to infauna gradients. Infauna gradients explained 37% of the sample distribution (PERMANOVA Adonis *p* < 0.01; Fig. 1G). Differential abundance analysis (MaAsLin2, *p* < 0.05, q < 0.09) showed 46 DNA virus bins with significant differences (Fig. 1F) (26 with *q* < 0.05; Supplementary Data 1). All DNA viruses, except *Autographiviridae*_386, increased in abundance in the HM, HMM and Baseline conditions relative to the LM condition. *Peduoviridae* viruses dominated the shift, with 18 viruses increasing in HM and HMM conditions, consistent with their overall dominance in the DNA virus community (59.6%).

#### RNA viruses

A total of 118 RNA virus contigs were obtained (Supplementary Data 2) and identified into 16 families (Fig. 1B; Table S1). Based on relative abundance, the community was dominated by *Yueviridae* (negative-sense ssRNA; 71.2%), followed by *Narnaviridae* (+ssRNA; 23.5%), and *Dicistroviridae* (positive-sense ssRNA; 4.9%), which are commonly associated with invertebrate, protist, and fungal hosts, respectively. Other RNA virus families, including *Tombusviridae*, *Phenuiviridae*, and *Spinareovirinae* were detected at very low relative abundances (0.01-0.0004%). The total abundance of RNA viruses did not differ among the infauna treatments (Fig. 1C), although the baseline samples showed significantly lower values. Similarly, diversity analyses revealed reduced RNA virus richness under HM compared to LM, with baseline samples exhibiting the lowest richness (Fig. 1E; Table S3).

PCoA showed significant compositional shifts (*p* < 0.05) among treatments (Fig. 1G) and explained 30% of the sample distribution. However, single RNA virus differential abundance (MaAsLin2, *p* < 0.05, *q* < 0.09) showed no significant differences between treatments (as abundance and diversity). This reflects differences detected at the community level rather than at the level of individual RNA virus abundances.

### Variables driving virus communities and co-abundance

We investigated the ecological drivers shaping viral community abundance and composition using multiple analytical approaches. The explanatory power of infauna varied depending on which bacterial proxy was used. When bacterial abundance from DNA metagenomes was the primary predictor (a proxy for bacterial biomass), infauna was not significant. However, when bacterial activity from RNA metatranscriptomes was used (a proxy for bacterial activity), infauna emerged as a significant predictor (Table S4-5).

When bacterial abundance from DNA metagenomes was used as the primary predictor, it dominated the regression models (Fig. 2A), showing a strong correlation with DNA virus abundance (McFadden’s R² = 0.96; *p* < 0.001) and explaining ∼19% of viral community compositional variation (PERMANOVA, *p* < 0.01; Fig. 2R; Table S6). In this scenario, infauna and other variables were not significant predictors (Fig. 2C-H), suggesting that DNA viruses simply track bacterial numbers in the sediment (bottom-up control). However, when bacterial activity from RNA metatranscriptomes (Fig. 2B) replaced bacterial abundance, infauna emerged as a significant predictor of DNA virus abundance (McFadden’s R² = 0.13; *p* < 0.001; Fig. 2C).

**Fig. 2.**
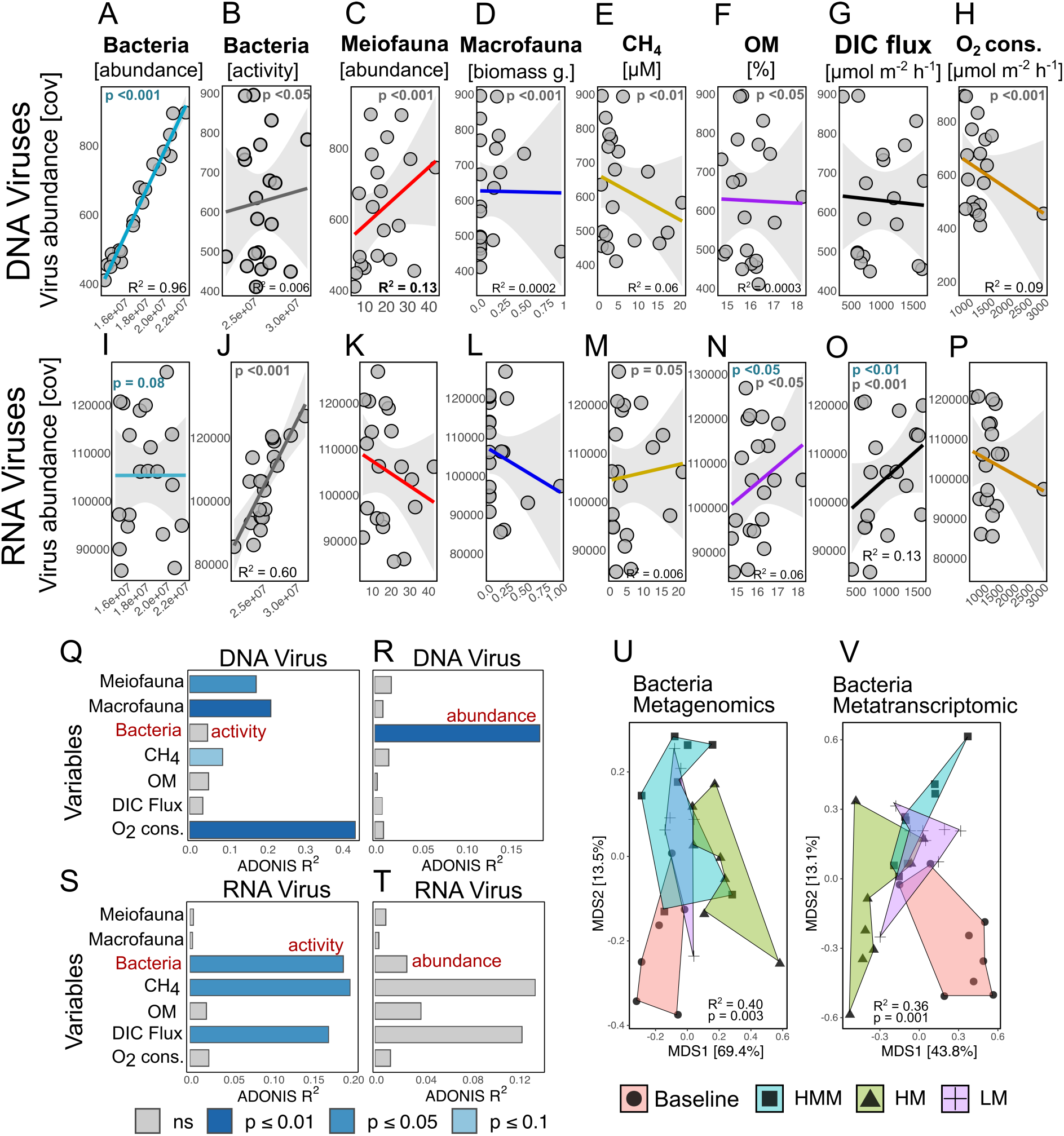
Environmental and biological drivers of viral community abundance and composition. Negative binomial generalised linear models (GLMs) for viral abundance are shown for DNA and RNA viruses using the following explanatory variables: (**A-P**) bacterial abundance and activity, meiofauna abundance, macrofauna biomass (g), CH_4_ concentration (µM), organic matter content (OM, %), dissolved inorganic carbon flux (DIC flux), and oxygen consumption (O_2_ cons.). McFadden’s R^2^ is shown when *p* < 0.05. (**Q-T**) Variance in viral community composition (R^2^, 0-1 scale) from PERMANOVA tests, with significance indicated by grey-blue shading, comparing models with bacterial abundance versus activity. (**U-V**) PCoA plots of bacterial abundance and activity based on relative abundance and Bray-Curtis dissimilarity (PERMANOVA, *p* < 0.05).

This shift is striking; infauna explained ∼18% of viral community composition (PERMANOVA *p* < 0.05; Fig. 2Q), and 40% with macrofauna biomass, while CH_4_ concentration, and O_2_ consumption are significant as well, while CH_4_ concentration, and O_2_ consumption. O_2_ consumption was the strongest individual predictor (R² = 44%; PERMANOVA p < 0.01; Fig. 2Q), followed by macrofauna (R² = 22%; PERMANOVA p < 0.01).

In contrast, RNA viruses showed no significant associations with meiofauna or macrofauna (Fig. 2K-L, 2T; Table S7-8). RNA virus abundance correlated with bacterial activity (McFadden’s R² = 0.60; *p* < 0.001; Fig. 2J), which explained 19% of compositional variation (PERMANOVA *p* < 0.05; Fig. 2S; Table S9). Additionally, OM% (Fig. 2N) and DIC flux (Fig. 2O) were significant explanatory variables. Notably, CH_4_ was significant (p <0.05; Fig. 2M) and relevant to community composition (R^2^ = 20%; PERMANOVA *p* < 0.05; Fig. 2S) only when bacterial activity was used as a variable.

Bacterial communities (Supplementary Data 2) clustered according to infauna gradients (Fig. 2U-V). To further investigate the relationship between DNA viruses and bacteria, we constructed co-abundance networks using SparCC (Fig. 3A). The resulting network (two-sided *p* ≤ 0.05) revealed unknown, *Zobellviridae*, *Peduoviridae*, and *Adenoviridae*-like viruses as central hubs, co-occurring with bacteria including *Gallionella*, *Desulfobulbus*, *Woeseia*, and *Leptothrix*. In addition, methanotrophic bacteria including *Methylomonas, Methylolococcus* and *Methylobacter* also showed co-abundance patterns with *Peduoviridae* and *Adenoviridae*_like viruses.

**Fig. 3.**
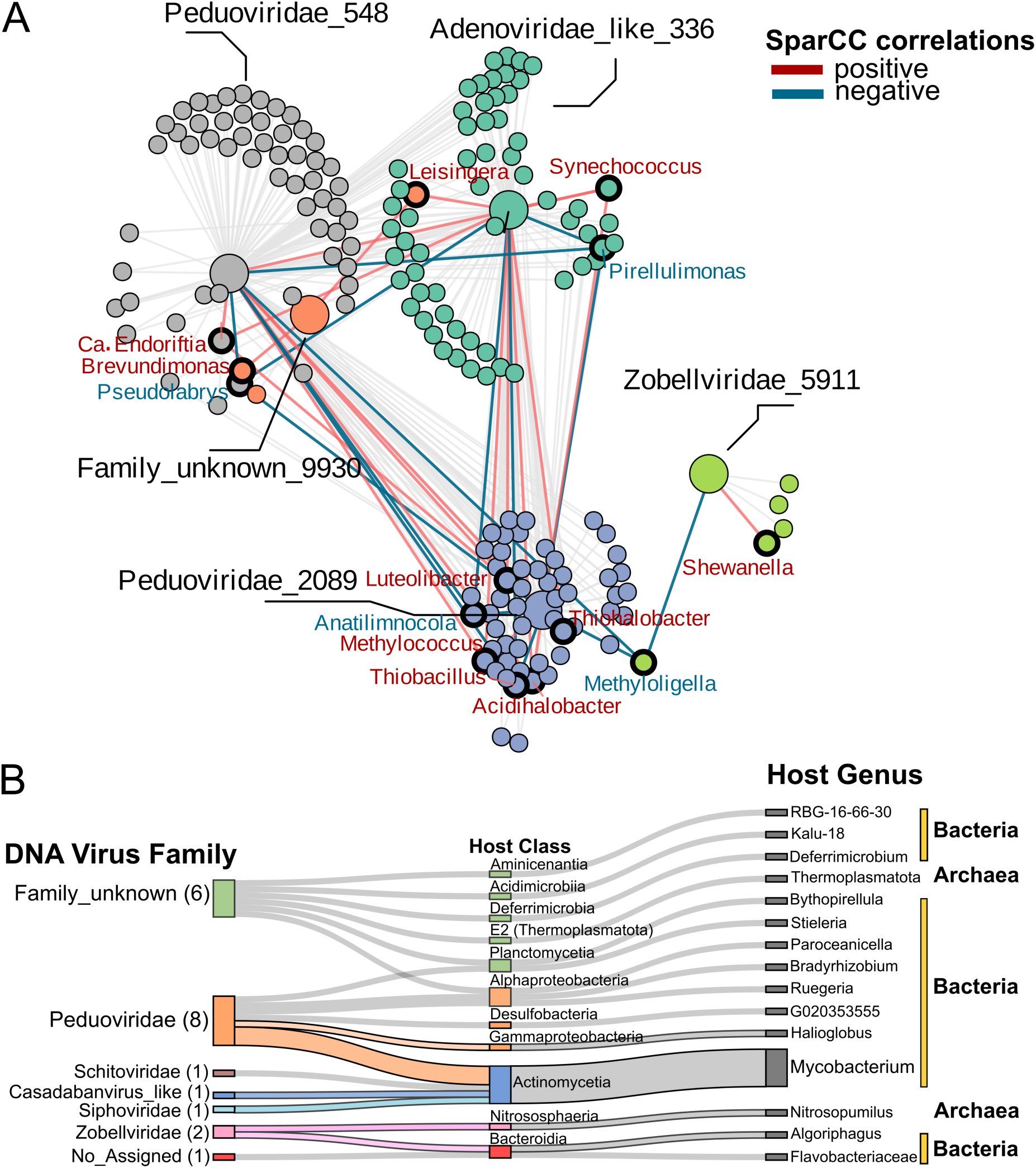
Co-abundance network and predicted hosts of DNA viruses. (**A**) SparCC co-abundance network of DNA viruses and bacterial taxa (two-sided *p* ≤ 0.05). Positive correlations are shown in red and negative correlations in blue; black edges highlight the strongest connections. Key bacterial genera are labelled. (**B**) Predicted host associations for DNA viruses (family level) inferred from sequence-based analysis. Viral genome counts are shown in parentheses, and ribbon colours indicate viruses with differential abundance across treatments (MaAsLin2, *p* < 0.05).

Host predictions based on sequence homology revealed *Peduoviridae* viruses linked to bacteria from several genera, including *Mycobacterium*, *Ruegeria*, *Bradyrhizobium*, and *Halioglobus* (Fig. 3B). Several unclassified DNA viruses were predicted to infect bacteria *Deferrimicrobium* and archaea within Thermoplasmatota. *Zobellviridae* viruses were predicted to infect organic matter degraders (*Algoriphagus*) and ammonia-oxidizing archaea (*Nitrosopumilus*).

### Infection cycle and activity of DNA viruses across infauna gradients

In terms of infection cycle, temperate viruses (57 %) were dominant, identified through the presence of integration/excision genes, compared to viruses potentially in the lytic cycle (43%), lacking temperate marker genes (Fig. 4A).

**Fig. 4.**
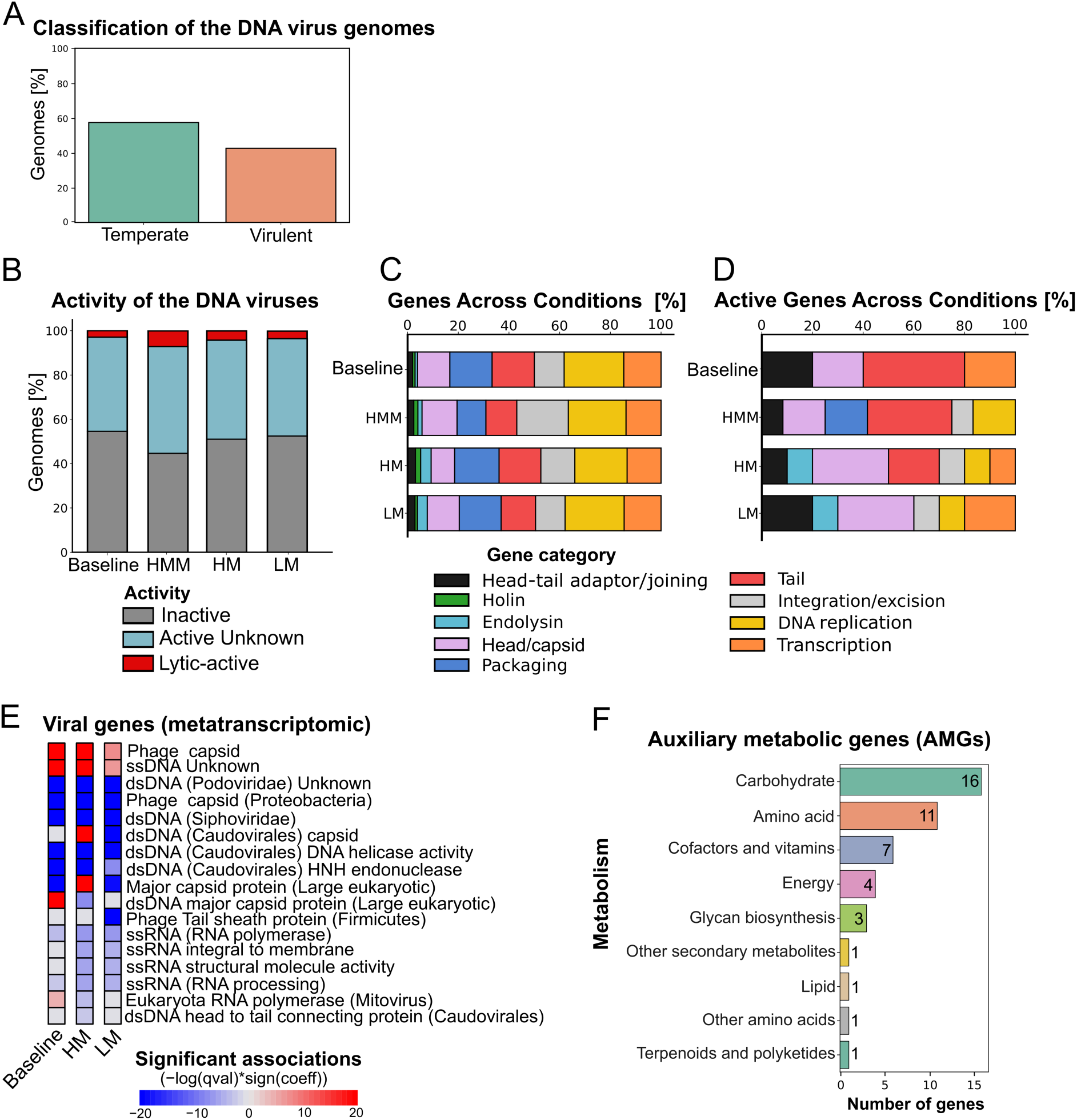
Infection cycles, transcriptional activity, and functional gene expression of DNA viruses. (**A**) Infection cycle types of DNA viruses inferred from the presence of temperate genes. (**B**) Percentage of DNA viral genomes classified as Inactive, Active Unknown and Lytic-active according to the functional annotation of expressed genes (adjusted average coverage >1×). (**C**) Relative abundance of lytic and lysogenic marker genes in DNA virus genomes across the treatments. (**D**) Relative abundance of the corresponding transcripts (adjusted average coverage >1×). (**E**) Differential expression (MaAsLin2, *q* < 0.05) of structural and replication genes (log-transformed adjusted coverage) grouped by gene type according to the RefSeq database. (**F**) Distribution of auxiliary metabolic genes (AMGs) across metabolic categories; the bar height indicates the gene count.

Transcriptional activity profiling showed that most DNA viruses remained transcriptionally inactive across treatments (60-68%) (Fig. 4B). Among the active viruses, the majority exhibited transcription of replication and structural genes (30-35%) without detectable expression of lysis genes, *e.g*., holin and endolysin, whereas viruses with detectable lysis gene expression represented a small fraction (≤5%) (Supplementary Data 3). The highest proportion of viruses with lysis gene transcription occurred under the HMM condition (high infauna + macrofauna), whereas the LM treatment showed the lowest overall viral activity.

Gene presence analysis revealed that DNA virus genomes were dominated by structural and replication genes (*e.g*., head, tail, packaging, and DNA polymerase) across all infauna treatments, whereas lysis-related genes were rare in terms of genomic representation (Fig. 4C), consistent with their low copy number (and late expression during the lytic cycle). Consistently, transcriptional activity was primarily associated with virion structure genes including head-tail adaptor, head, tail, among others, and replication machinery (Fig. 4D Fig S2). This expression pattern was further supported by differential gene expression analysis (MaAsLin2, *p* < 0.05), which showed significantly higher expression of viral structural genes under the HMM condition than under the Baseline, HM, and LM treatments (Fig. 4E). While DNA viral genes dominated the transcriptional response, a minor but significant increase in the expression of specific ssRNA viral genes was also detected, including RNA-dependent RNA polymerases and structural proteins associated with mitoviruses and other eukaryotic-infecting lineages.

A small yet functionally diverse set of auxiliary metabolic genes (AMGs) was identified in DNA virus genomes (Fig. 4F), primarily linked to carbohydrate and amino acid metabolism, energy production, and cell envelope biosynthesis (Fig. S3; Supplementary Data 3). Nevertheless, mapping of metatranscriptomic reads against these gene sequences yielded negligible coverage, detecting no activity of those genes across the infauna treatment.

## Discussion

### DNA and RNA viruses in Storfjorden sediments

Our results underscore the ecological importance of lytic dsDNA phages in the Storfjorden sediments as evidenced by the recovery of DNA viruses across 21 families and dominated by *Peduoviridae*. Non-phage *Adenoviridae*-like viruses were also abundant (∼3%), whereas the large fraction of unclassified DNA viruses highlights the extent of viral “dark matter” (Paez-Espino et al. 2016; Wolf et al. 2020) and unexplored genomic diversity within benthic viral assemblages. Comparable patterns of dsDNA phage dominance have been observed in coastal and deep-sea sediments (Yu et al. 2024), suggesting that phage-driven microbial turnover is widespread in sedimentary ecosystems.

The study of marine RNA viruses is relatively recent but our understanding of their diversity has greatly expanded with meta-omics advances and the application of AI-assisted sequence annotation (Zayed et al., 2022; Liao et al., 2022; Hou et al., 2024). RNA viruses from this study belong to 16 families (Fig. 1B) with eukaryotic viruses *Yueviridae, Narnaviridae*, and *Dicistroviridae* being the most abundant. As members of these families are associated with invertebrates, protists, and fungi (Valles et al., 2017; Charon et al., 2021; Wu et al., 2024), it indicates strong connections between RNA viruses and eukaryotic hosts. While some RNA viruses appeared highly abundant, consistent with reports that RNA viruses may be equal to or even exceed DNA viruses in marine systems (Culley et al., 2014; Urayama et al., 2018; Vlok et al., 2019), only DNA viruses showed significant shifts in structure, abundance and diversity across the infauna gradients, suggesting different community drivers for RNA and DNA viruses in sediments.

### Infauna**-**driven enhancement of DNA viral abundance and diversity

We found that high abundances of infauna correlated with an increased abundance and diversity of DNA viruses and altered community composition. In particular, *Peduoviridae* increased in abundance, supporting previous reports that infauna influence viral and microbial distributions in fjords (Wrobel et al., 2013). DNA viral abundance was strongly correlated with metagenomic bacterial abundance (Fig. 2A), whereas meiofauna, macrofauna and O_2_ consumption arose as significant explanatory variables when replaced with metatranscriptomic bacterial activity (Fig. 2B). Furthermore, bacterial communities clustered according to infauna gradients (Fig. 2U-V), suggesting that infauna shape sediment structure, organic matter availability, and bacterial activity. Together, these results suggest that sediment mixing, solute transport, and particle resuspension by infauna influence DNA viral communities through two interconnected mechanisms. First, infauna modulate bacterial abundance and activity (bottom-up control). Second, because viruses are non-motile and depend critically on spatial proximity to hosts (Meysman et al., 2006), infaunal bioturbation likely enhances virus-host encounter rates (top-down control) by disrupting sediment structures and generating pore-water flows that move particles, including bacteria and viruses. This increased contact between viruses and diverse bacterial hosts would be expected to amplify infection dynamics, particularly in metabolically active bacteria.

The microhabitats created by burrow ventilation provide a substrate for this encounter-enhanced infection process. Burrow ventilation enhances organic carbon, oxygen, and nitrate delivery, stimulating metabolically active copiotrophic bacteria (Kristensen et al., 2005; Bertics & Wiebke, 2009) and creating oxic-anoxic interfaces with microscale chemical heterogeneities. This environmental patchiness diversifies and expands the niche space, promoting microbial biodiversity (heterogeneity-begets-diversity; Levin, 1992; Glud 2008). Critically, the increased encounters between viruses and this diverse bacterial community, enabled by enhanced mixing, allow viral infection of bacteria occupying contrasting redox niches. This selective pressure drives the coexistence of both lytic and temperate viral strategies adapted to different host metabolic states and redox conditions (Howard-Varona et al., 2017). These dynamic microhabitats accelerate microbial turnover and virus-host interactions, reinforcing viral-host coevolution. For instance, giant multicellular bacteria forming mats under high phage pressure harbor expanded antiviral defence systems compared to unicellular taxa (Fonseca et al., 2025), illustrating how the combination of enhanced encounter rates and microbial diversity under bioturbation promotes ecological and evolutionary coupling between viruses and their bacterial hosts in sediments. Thus, our data indicate that infauna can act as central ecological modulators of sedimentary DNA viral communities through intertwined physical, biogeochemical, and biological pathways

### Divergent responses of RNA viruses

RNA viral communities exhibited compositional shifts across infauna treatments (Fig. 1G), but these changes were not accompanied by differences in the total abundance, alpha diversity, or individual viral taxa. This suggests that infauna disturbance restructures RNA viral assemblages through neutral species turnover, likely driven by environmental shifts, rather than promoting the broad-scale amplification or dominance of specific lineages observed in DNA viruses. While DNA viruses respond to the direct metabolic ‘engine’ of bacterial hosts, the RNA virome likely undergoes passive rebalancing as the physical and chemical structure of the sediment changes. This decoupling reflects the distinct host spectra and life-history traits of RNA viruses, which primarily infect eukaryotic hosts (Zhang et al. 2024), including protists, fungi, and multicellular animals (Urayama et al. 2018). Their populations are more sensitive to organic matter quality and stable redox structures than to small-scale sediment reworking. Additionally, their general short environmental persistence, high mutation rates, and lack of a lysogenic phase (Sime-Ngando, 2014; Gustavsen et al., 2014) further decouple them from the ecological feedback that strongly links DNA viruses to bacterial hosts.

RNA viruses showed stronger correlations with bacterial activity, DIC fluxes, and CH₄ (Fig. 2S). We interpret these results as reflecting a shared sensitivity to current environmental conditions rather than a direct response to infauna-driven disturbance. Specifically, RNA viruses track the same abiotic ‘chemical stage’ (O₂, CH₄, organic matter redistribution) as bacterial communities, but this co-occurrence is decoupled from the direct mechanical effects of bioturbation that amplify DNA viral abundance. This interpretation is supported by the absence of responding RNA phages and the dominance of eukaryotic-associated families: *Yueviridae* and *Narnaviridae* (crustacean hosts; Kibenge, 2024), and *Dicistroviridae* (fungal hosts; Starr et al., 2019). Their hosts likely respond at temporal and spatial scales that differ from those of bacteria. Thus, this correlation may reflect synchronous responses to the same environmental gradient rather than a mechanistic response to bioturbation.

While bioturbation modifies sediment oxygenation and organic matter redistribution, these shifts may not immediately translate into changes in viral host availability within a 10-day experimental timeframe. An important difference lies in host size and temporal responsiveness: bacterial populations respond metabolically within hours to reworked oxygen and nutrient pulses (Luna et al., 2002) and are substantially smaller (0.5-5 μm) than their eukaryotic counterparts. Because bacteria are small, they are preferentially mobilized by infauna burrowing and ventilation, which generates pore-water flows that resuspend small particles, whereas larger eukaryotic organisms remain relatively stationary. This size-dependent differential mobility may substantially enhance the encounter rates between viruses and their small bacterial hosts. In contrast, macrofaunal invertebrates (e.g., crustaceans associated with *Yueviridae*) that serve as RNA virus hosts have generation times of months to years, and their larger body size means they are not preferentially mobilized by bioturbation. Consequently, the selective response of DNA viruses, but not RNA viruses, to bioturbation reflects the differential efficiency with which meiofaunal disturbance mobilizes small bacterial hosts compared to larger eukaryotic organisms, combined with the temporal mismatch between rapid microbial responses and slower host population dynamics. While both viral groups experience identical physical reworking, their divergent responses appear driven by the interaction of host size, metabolic responsiveness, and population-level demographic timescales

### Infauna-bioturbation reshapes viral infection strategies

Lytic-active viruses dominated the high-infauna treatments, consistent with enhanced virus-host encounter rates under intensified sediment mixing. In contrast, temperate viruses were similarly transcriptionally active across the infauna treatments. These patterns indicate that an increase in infauna abundance promotes both viral activation and shifts toward lytic infection strategies. Because viruses are non-motile, their dispersal and host contact rates depend strongly on sediment reworking and pore water transport (Meysman et al., 2006). In addition, infauna burrowing and irrigation generate dynamic redox and physicochemical gradients that may act as stressors to promote prophage induction within localized microhabitats (Howard-Varona et al., 2017).

Across all infauna levels, viral transcription was dominated by genes involved in genome replication and virion assembly processes. In lytic bacteriophages, genes encoding capsids, tails, and lysis functions are typically expressed during the late stages of infection, following or overlapping with viral genome replication, reflecting the conserved temporal regulation of phage transcription (Blasdel et al., 2017). The concurrent expression of replication, and structure-related genes observed here likely reflects asynchronous infections occurring across diverse microbial hosts, consistent with sustained virion production rather than synchronized infection.

Auxiliary metabolic genes (AMGs) identified in viral genomes are primarily associated with carbohydrate metabolism, amino acid metabolism, and central energy-related pathways. These functions are commonly reported in environmental viral genomes and are thought to support host metabolic processes during infection rather than introducing novel pathways (Breitbart et al., 2018; Coutinho et al., 2018). In this context, the detected AMGs represent the viral genomic potential to modulate host metabolic fluxes during infection without implying measurable changes in host physiology. Similar AMGs have been linked to processes such as organic matter degradation and substrate utilization in marine sediments, including chitin-associated pathways (Middelboe et al., 2025), suggesting that viral infections may influence the turnover of labile organic compounds under bioturbation conditions.

From an evolutionary perspective, physical mixing and increased microbial contact rates are key determinants of the efficiency of horizontal gene transfer (Brochier-Armanet & Moreira, 2014). By intensifying virus-host interactions and infection frequency, infaunal bioturbation may enhance the opportunities for viral-mediated gene exchange, even if such metabolic modulations are transient or niche-specific, within sediment microbial communities. This adds an evolutionary dimension to sediment mixing, positioning viruses not only as agents of microbial mortality but also as dynamic components of faunal-microbial-geochemical feedback in bioturbated sediments.

### Ecological implications

Our results indicate that infauna act as a structural driver of sedimentary DNA viral communities, advancing a view of benthic ecosystems in which faunal, microbial, and viral dynamics are ecologically coupled rather than operating in isolation. (Fig. 5). While these findings, based on a 10-day incubation in a representative brackish-water ecosystem, provide a high-resolution snapshot of this coupling, we acknowledge that the strength of these interactions may vary across benthic habitats and temporal scales. For instance, a 10-day window primarily captures the immediate responses of fast-responding microbial and viral lineages. Longer-term studies might reveal delayed successional shifts, particularly for the RNA virome and its eukaryotic hosts, whereas shorter pulses might capture transient lytic events that occur within days of the initial disturbance. Nevertheless, our data extends the previous work from the same experimental system, showing that infauna regulate methane dynamics primarily through physical transport and bacterial activity (Broman et al., 2024). Here, we show that viral communities respond to the same infauna-associated disturbance. This suggests that viruses may represent an additional, previously overlooked component that links faunal activity to microbial turnover in bioturbated sediments.

**Fig. 5.**
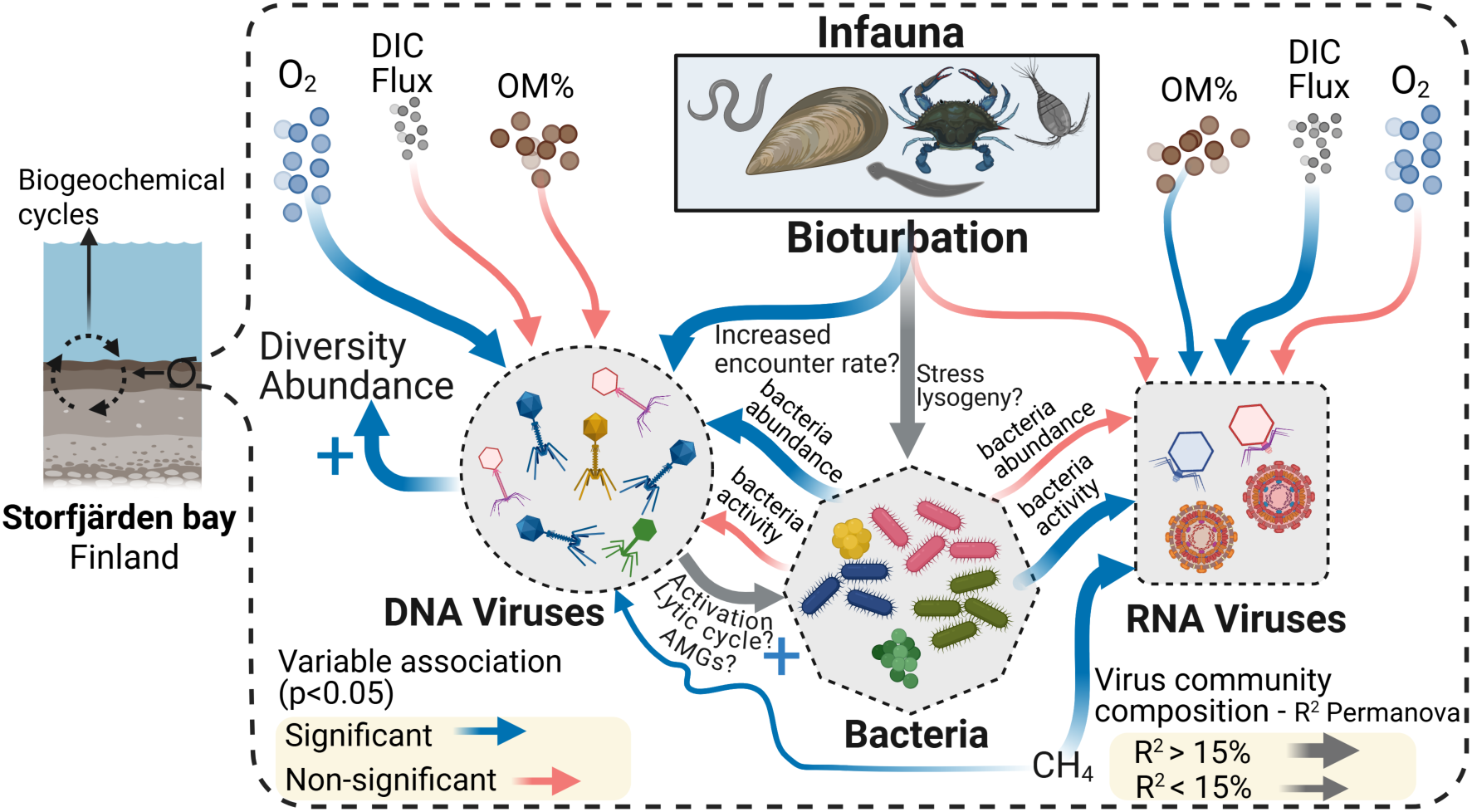
Conceptual model of infauna-virus interactions in sediments. Schematic overview of the relationships between abiotic variables (CH_4_, organic matter content [OM%], dissolved inorganic carbon flux [DIC flux], and oxygen consumption [O_2_]) and biotic components (infauna: meiofauna and macrofauna, and bacteria) with DNA and RNA viruses. Blue arrows indicate significant relationships (*p* < 0.05) identified by regression models (Fig. 2A-P), whereas the red arrows denote non-significant relationships (*p* > 0.05). The arrow width reflects the proportion of viral community variation explained by each variable based on PERMANOVA (Fig. 2Q-T): wide arrows represent >15% of the explained variance; thin arrows represent <15% or non-significant effects (*p* > 0.05). Grey arrows depict inferred direct or indirect relationships, and blue plus symbols denote positive effects. Dashed lines and question marks indicate hypothesized physical mechanisms (e.g., enhanced virus-host encounter rates via bioturbation).

In other sedimentary environments, viral infections of methanogens, methanotrophs, sulfate reducers, and iron-cycling bacteria have been shown to influence carbon and redox-sensitive element cycling through host mortality and metabolic modulation (Chen et al., 2020; Cheng et al., 2022; Zhong et al., 2024). Although such processes were not directly quantified here, the virus-infauna-microbe associations observed in this study are consistent with viruses acting as responsive components of the benthic biogeochemical feedback. Direct measurements of infection rates, host-virus linkages, and metabolite fluxes will help further resolve these interactions, particularly at short temporal scales following infauna disturbance.

In conclusion, higher infauna abundance was associated with increase in abundance, diversity, and transcriptional activity of DNA viruses, whereas RNA viral communities showed limited quantitative responses. These patterns indicate that infauna-mediated sediment reworking selectively structure DNA viral communities and suggest that alters the ecological context of virus-host interactions, potentially enhancing the coupling between bacteria and their viruses and accelerating the lytic cycle. By demonstrating that infauna gradients shape DNA viral community structure independently of direct physicochemical controls, this study opens a new dimension in our understanding of how faunal biodiversity may regulate microbial processes in coastal sediments.

## Methods

### Sediment samples and infauna incubations

Sediment cores (n = 63; inner Ø 4.6 cm, length 30 cm) were collected from Storfjärden Bay, Finland (59.8559°N, 23.26695°E; 34 m depth) using a box corer in May 2022. Each core contained ∼14 cm of sediment and ∼14 cm of water. The in-situ conditions were 6.5°C, 11.9 mg O_2_ L^-1^ and 6.4 PSU. Three sediment cores were sliced onboard (1 cm intervals) to determine porosity and organic matter (OM %) using loss on ignition (LOI).

Sediment cores were used to establish five experimental infauna treatments (*n* = 12 independent sediment cores for each treatment): low meiofauna (LM), low meiofauna + macrofauna (LMM), high meiofauna (HM), high meiofauna + macrofauna (HMM), and unmanipulated sediment cores (Baseline). Importantly, the present study (metagenome sequencing) considered LM, HM, HMM, and Baseline. To set up the treatments, meiofauna were extracted from 60 cores by the density separation method described in Nascimento et al. (2012) using a Levasil silica gel colloidal dispersion (1.21 kg m^-3^) following sequential sieving (1000 and 40 µm three times) and anesthesia with MgCl₂ (740 mM, 5 min). After the third isolation, the sieve was rinsed with seawater, and meiofauna from the two cores were pooled and returned to one core (HM), while the second core received no meiofauna and was designated as LM. Fine sediment particles passing through the 40 µm sieve were returned to both cores. This resulted in the following meiofauna abundance: average abundances (×10^3^ m^-^_2_): LM = 46 ± 16, HM = 173 ± 65, HMM = 214 ± 111, and Baseline = 261 ± 100. Additional sediment was sieved to collect *Macoma balthica* (10-12 mm), and two individuals were added to half of the high meiofauna abundance cores to obtain densities comparable to those found in natural conditions (1200 individuals m^−2^. Gogina et al., 2016). Overall, macrofauna abundance was therefore: LM 0 ± 0, HM 0 ± 0, HMM 2 ± 1, and Baseline 1 ± 1 (average ± SD per treatment, sediment core surface area: 16.6 cm^−2^). The fauna-treatment sediment cores were kept in aerated in situ bottom water (∼6°C) with controlled stirring and acclimated for 10 days in the dark. Oxygen, temperature, and salinity levels were monitored daily to maintain stable conditions.

### Termination and infauna quantification

Ten days after the addition of the infauna and after 6 hours of incubation for flux measurements (CH4 and DIC sediment-water flux, as well as the O_2_ flux, i.e., oxygen consumption), the experiment was terminated, and the infauna was quantified (Fig. 6A-B).

**Fig. 6.**
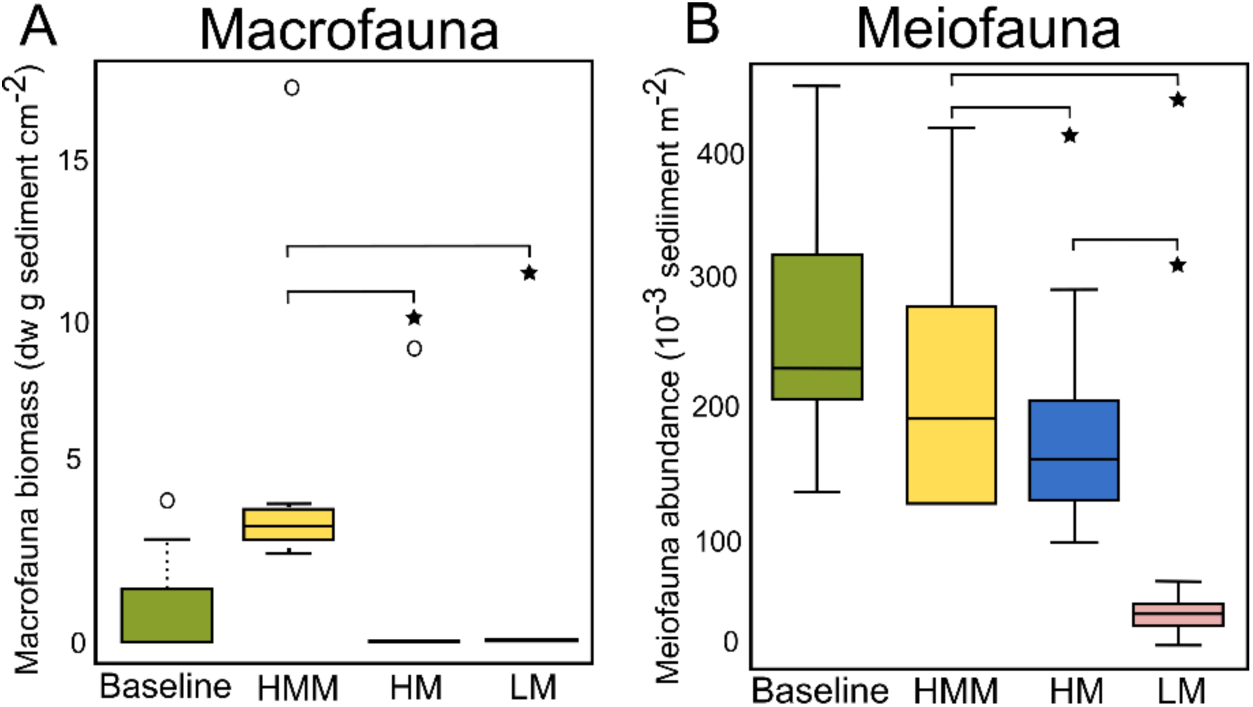
Biomass of macrofauna and abundance of meiofauna under different treatments. Sediment cores were experimentally adjusted for meiofauna and macrofauna and acclimated for 10 days. The treatments comprised HM = high meiofauna, HMM = high meiofauna + macrofauna, LM = low meiofauna, and unmanipulated cores (Baseline). (**A**) Macrofaunal biomass measured from all cores (0-14 cm depth, n = 7 per treatment). dw = dry weight. (**B**) Meiofaunal abundance determined from the uppermost sediment layer (0-1 cm, n = 7 per treatment). In the boxplots, the horizontal line indicates the median, the boxes represent the interquartile range, and the whiskers extend to the smallest and largest values. The circles denote outliers ≥1.5 × the box length. Start depicts Benjamini-Hochberg p-adjustment, *p* < 0.05. Baseline cores showed meiofauna and macrofauna levels within the treatment range, confirming in situ realism.

Macrofauna from all sediment cores were retrieved via sieving (1000 µm), taxonomically identified, enumerated, and their biomass estimated from intact individuals based on dry-weight loss after oven-drying at 60 °C for 24 h. Meiofauna were extracted from frozen sediment subsamples (2-8 mL) collected from the 0-1 cm sediment layer of each replicate (see Table S10 for taxonomy), following thawing, using a density-based separation procedure described previously. Extracted meiofauna were subsequently sorted, identified, and counted under a stereomicroscope at 50× magnification.

### Chemical analyses

After the termination of the experiment, three cores per treatment were sliced at 1 cm intervals for porewater CH_4_ analyses, while 7 cores by treatment (except HMM, *n* = 6) were used for sediment-water flux (6 h) of CH_4_ and DIC, as well as the O_2_ (oxygen consumption) (Supplementary Data 4). DIC flux for each treatment was measured by filtering water through a 0.45 μM glass fiber filter into 12 mL Exetainer glass vials and stored in the dark until analysis. Oxygen consumption rate measured in the water phase on top the sediment surface in each core was measured by inserting a microelectrode (OX-, Unisense) before and after the incubations. Porosity and OM% were determined from dried samples (105 °C, 24 h) and LOI (550 °C, 4 h) measurements (procedures are described in detail by Broman et al. [2024]).

### DNA and RNA extraction and sequencing

DNA and RNA were extracted from the 0-1 cm surface sediment (∼2 g) using the DNeasy PowerSoil Pro and RNeasy PowerSoil Total Kits (Qiagen). To obtain RNA, the DNA was removed using the TURBO DNA-free kit from Invitrogen, and library preparation was carried out with the TruSeq Stranded mRNA Kit from Illumina. RNA libraries were prepared from 31 cores (HMM: 8, HM: 8, LM: 7, and Baseline: 8), and DNA libraries were prepared from 26 cores (HMM: 7, HM: 7, LM: 6, and Baseline: 6; Supplementary Data 4)), both using Illumina DNA PCR-free. DNA and RNA libraries were sequenced on an Illumina NovaSeq 6000 (2 × 150 base pairs).

### Bioinformatic analyses

#### DNA and RNA library quality control and pre**-**processing

Reads were quality controlled with FastQC 0.11.9 (Andrews, 2010) and MultiQC 1.12 (Ewels et al., 2016) before and after trimming. Trimming involved removal of Illumina adapters using SeqPrep 1.2 with default settings aimed at adapter sequences (St John, 2011). PhiX control sequences were then removed following alignment to the PhiX genome (NCBI Reference Sequence: NC_001422.1) using bowtie2 2.5.2 (Langmead and Salzberg, 2012). Low-quality and short reads were filtered using Trimmomatic 0.39 (Bolger et al., 2014) with the parameters LEADING:20, TRAILING:20, and MINLEN:80. The resulting quality-controlled DNA libraries (*n* = 27) averaged 101 million paired-end reads and RNA libraries (*n* = 31) averaged 71.4 million paired-end reads.

#### DNA virus identification

Quality-trimmed metagenomic reads were co-assembled using MEGAHIT v1.2.9 (Li et al., 2015) with default parameters. The resulting contigs were binned using SemiBin2 Version 2.1.0 (Pan et al., 2022) with the mode “single_easy_bin” and -minfasta-kbs 5 with alignment files generated by mapping trimmed reads to contigs using Bowtie2 v2.5.2 and SAMtools v1.20 (Li et al., 2009). Bins up to 1.2 Mb in size were selected to target bacteriophages and viruses that range from ∼2 kb to over 1 Mb. Size selected bins were and used as input to PHAMB (Johansen et al., 2022) which uses a random forest classifier to discern the viral bins (Fig. 7) and includes DeepVirFinder (Ren et al., 2020) to predict viral sequences. PHAMB was used with the Virus Orthologous Groups Database (VOGDB., June 2024) that includes all virus genomes from RefSeq. The viral bins were evaluated using CheckV v1.0.3 (Naifach et al., 2021), and Medium- and High-quality bins with contamination score 0, and at least one viral gene was retained.

**Fig. 7.**
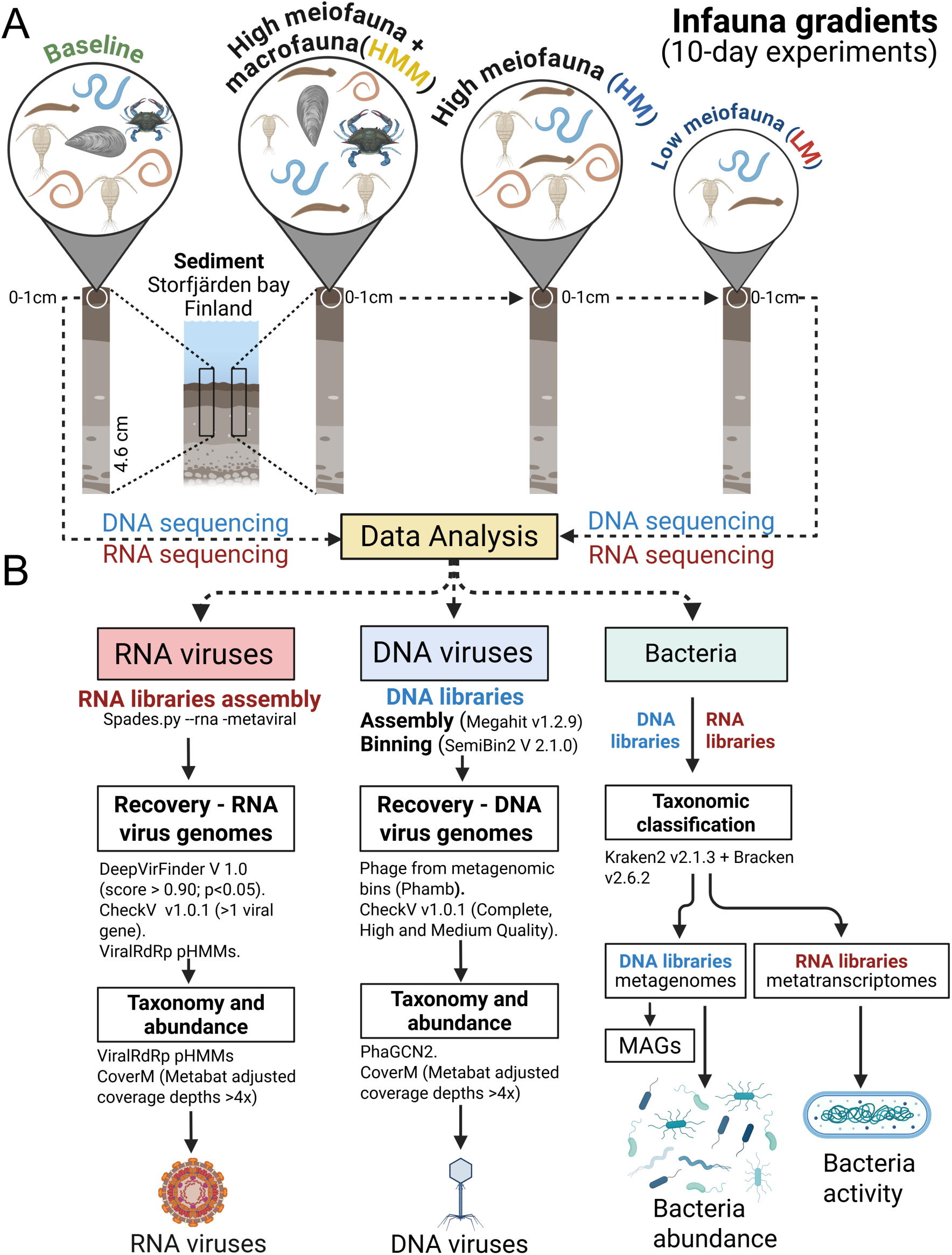
Experimental design and analytical workflow of the study. Infauna treatments included HM = high meiofauna, HMM = high meiofauna + macrofauna, LM = low meiofauna, and unmanipulated cores (Baseline). After 10 days, the upper 0-1 cm sediment layer was sampled for infauna quantification and for DNA and RNA extraction. RNA libraries were used to characterise RNA viruses, transcriptional activity, and bacterial activity, whereas DNA libraries were used to identify DNA viruses and assess bacterial abundance.

DNA virus bins were quantified by mapping the quality-filtered reads against them using Bowtie2 with the default parameters. The resulting bam files were used to estimate the viral abundance using CoverM v0.7.0 (Aroney et al., 2025) in “contig” mode with the “MetaBAT2 Adjusted Coverage Depths” method which adjusts for genome-wide coverage variability. To reduce noise and artifacts, a coverage threshold of 4x was applied, whereby viral genomes with a mean coverage below this value were set to zero in the abundance final matrix and normalized for downstream analysis. In total, 141 DNA virus genomes were obtained, including 3 complete genomes (Supplementary Data 1).

#### Identification of RNA Viruses

RNA viruses were identified from contigs generated by de novo assembly (single) of quality-trimmed metatranscriptomic reads using rnaSPAdes v3.15.5 (Fig. 7). Only contigs > 500 bp were retained for downstream analysis.

Putative RNA viral contigs were first identified using DeepVirFinder v1.0, applying a stringent threshold (score ≥ 0.90, p < 0.01). Redundant viral contigs were removed using CD-HIT-est v4.8.1 (Li and Godzik, 2006) at 100% nucleotide identity (-T 10 -c 1 -n 10 -M 8000). Open reading frames were predicted on the non-redundant contigs using Prodigal v2.6.3 (Hyatt et al., 2010) in the metagenomic mode (“-p meta -g 11 -T 8”). To further confirm RNA virus identity, RNA-dependent RNA polymerase (RdRp), a universal hallmark protein of RNA viruses, was detected using a curated set of RdRp profile hidden Markov models (pHMMs; https://github.com/ingridole/ViralRdRp_pHMMs), comprising 77 family level HMMs with HMMER v3.4 (Potter et al., 2018).

Final RNA virus contigs were required to meet all of the following criteria: (i) encoding a single RdRp gene, (ii) a DeepVirFinder score ≥ 0.90 (p < 0.01), and (iii) the presence of at least one viral gene as assessed by CheckV. Using these criteria, 348 RNA viral contigs were obtained. These contigs were subsequently quantified using the same mapping and normalization parameters applied to the DNA viruses.

#### DNA and RNA Virus Taxonomy

DNA virus taxonomy was determined using PhaGCN 2.0 program (Jiang et al., 2023), which uses a semi-supervised learning model for phage taxonomic classification. RNA virus taxonomy was determined from the RdRp signature analysis using the 77 family-level pHMMs described above. Both methods report taxa at the family level.

#### Classification of the infection cycle and activity of DNA viruses

To assess infection strategies and the transcriptional activity of DNA viruses, all curated viral bins were annotated using Prokka v1.14.5 (Seemann, 2014) with the VOGDB protein database (release 224) and eggNOG-mapper v2.1.12 (Cantalapiedra et al., 2021). Genomes were first screened for marker genes indicative of lysogeny (e.g., integrase, recombinase, excisionase, and prophage) or lytic/structural functions (e.g., capsid, tail, terminase, holin, and endolysin). Genomes containing at least one lysogeny-related gene were classified as temperate, reflecting the potential for lysogenic infection cycles. Viruses lacking such markers were classified as virulent, representing strictly lytic viruses.

Quality-controlled reads from metatranscriptomes were then mapped to the viral gene catalogue using Bowtie2 (--very-sensitive), and gene expression was quantified with CoverM using the “MetaBAT2 Adjusted Coverage Depths” mode, accounting for edge trimming and high-identity alignments. Gene-level activity was defined as an adjusted mean coverage of >1×, corresponding to detectable transcription. To determine viral genome activity, gene expression profiles were integrated with functional annotations. Structural and lytic genes were used as indicators of lytic activation. While this is an imitated diagnosis, it was a practical approach. Based on these profiles, each viral genome bin was assigned to one of three activity states: i) Inactive, no genes expressed above the threshold; ii) Active unknown, expression limited to non-structural or non-lytic genes; and iii) Lytic-active (actively replicating), at least one structural or lytic gene expressed above the threshold level.

Differential expression of viral genes (RefSeq database), including structural and replication genes (for example, RNA polymerase and DNA helicase), was performed using MaAsLin2 (*q* < 0.05) and log-transformed mode of adjusted coverage values. Finally, auxiliary metabolic genes (AMGs) were identified using Vibrant 1.2.1 (Kieft et al., 2020), and potential viral hosts were predicted using iPHoP v1.4.1 (Roux et al., 2023).

#### Bacterial taxonomy, abundance and MAGs

Bacterial abundance and taxonomy were determined from metagenomic reads using Kraken 2.0.9 (Wood et al., 2019) with the SILVA database, and Bracken v2.6.2 (Lu et al., 2017) was used to refine the genus-level assignments. The results were processed using Kraken-Biom and Biom software. The same objective was achieved with metatranscriptomic reads, as a proxy for bacterial activity, following Broman et al. (2024). Briefly, 16S SSU rRNA reads from metatranscriptomic libraries were extracted using SortMeRNA (Kopylova et al., 2012), classified using Kraken 2.0.9, and quantified using Bracken v2.6.2. Both data matrices were normalized to relative abundance and analyzed using the Phyloseq package (Mcmurdie and Holmes, 2013) in R v4. In addition, metagenome-assembled genomes (MAGs) of bacteria and archaea were obtained to investigate the potential hosts of DNA viruses (see Supplementary Data 2).

#### Diversity and Statistical Analyses

Viral diversity indices (Shannon, Simpson, and richness) were calculated using the vegan package (Dixon, 2003) in R v4.4.0. Total DNA and RNA viral abundance, based on cumulative MetaBAT2 adjusted coverage depths, and diversity indices were compared among treatments using Student’s t-tests and Wilcoxon rank-sum tests (*p* < 0.05). The differential abundance of individual viral genomes across infauna conditions was assessed using MaAsLin2 (Mallick et al., 2021) with linear models (LM), minimum prevalence = 0.05, log transformation, Benjamini-Hochberg correction, and significance thresholds of *p* < 0.05 and *q* < 0.09.

Community composition patterns were evaluated using Bray-Curtis dissimilarities and visualized using Principal Coordinate Analysis (PCoA). Group differences were tested using ANOSIM (2000 permutations) and PERMANOVA (Adonis, 999 permutations; *p* < 0.05) in vegan.

To identify environmental and biological predictors of viral abundance, we compared Generalized Linear Models (GLMs) with Gaussian (LM), Poisson, and Negative Binomial error distributions, as well as Generalized Additive Models (GAMs), using AIC to select the best-fitting model. The final Negative Binomial GLM (MASS v7.3-65) included bacterial abundance and activity, meiofauna abundance, macrofauna biomass, CH_4_ concentration, OM%, DIC flux, and O₂ consumption as explanatory variables. Overdispersion, multicollinearity, and goodness-of-fit (McFadden’s R², null, and residual deviance) were evaluated, and variable significance was tested using the Wald test. The relationships between viral community composition and explanatory variables were further explored using Correspondence Analysis and Redundancy Analysis (RDA) using vegan and ggplot2. Co-occurrence networks between DNA viruses and bacteria were inferred using SparCC (Friedman & Alm, 2012) with 200 bootstrap samples (two-sided *p* < 0.05).

## Acknowledgments

We thank Elias Broman for setting the infauna gradient experiments, Laura Kauppi for the support in the field plus laboratory, and Jaana Koistinen for the support in the laboratory. We acknowledge the Centre for Coastal Ecosystem and Climate Research (jointly operated by University of Helsinki and Stockholm University) for institutional support. Sequencing data were generated with assistance from the National Genomics Infrastructure in Stockholm, supported by Science for Life Laboratory, the Knut and Alice Wallenberg Foundation, and the Swedish Research Council. Computational resources and support were provided by NAISS/Uppsala Multidisciplinary Center for Advanced Computational Science. Bioinformatic analyses were carried out using NAISS-allocated computational resources (projects 2023/22-1280 and 2024/22-1008) at UPPMAX through the Swedish National Infrastructure for Computing (SNIC).

## Funding

Swedish Research Council grants to FJAN (grant no. 2024-04627) and KH (grant no. 2022-04340). Marcus and Amalia Wallenberg Foundation. Stockholm University internal project number: 31004348 (FJAN). The work was supported by the Danish National Research Foundation through the Center for Hadal Research (DNRF145) to MM.

## Author contributions

Conceptualization: AF, FJN

Methodology: AF, KH, FJN, MM, EM

Investigation: AF, FJN, KH

Visualization: AF

Supervision: FJN, MM, KH

Writing original draft: AF

Writing, review & editing: AF, FJN, KH, MM, EM, CH, AN

## Competing interests

Authors declare that they have no competing interests.

## Data and materials availability

All data are available in the main text or in the supplementary materials, including supplementary figures and tables in the Supplementary Material file. Supplementary data with extensive tables are provided in xlm format: Supplementary Data 1, Data 2, Data 3, and Data 4. Raw DNA and RNA sequencing data are available in the NCBI repository under BioProjects PRJNA1365460 and PRJNA917468, respectively.

## Notes

### Competing Interest Statement

The authors have declared no competing interest.

