## Supplementary_Materials for "Infauna selectively enhance DNA virus diversity and activity in marine sediments"

### Infauna shape DNA viruses in sediments

**This PDF file includes:**

Tables S1 to S10
Figs. S1 to S3

**Table S1.** DNA and RNA virus family abundance (%).

| DNA virus Family | Abundance (%) |
|---|---|
| Peduoviridae | 54.58 |
| Family_unknown | 22.81 |
| Adenoviridae_like | 7.2 |
| Siphoviridae | 3.43 |
| Inoviridae_like | 2.81 |
| Zobellviridae | 1.82 |
| Gracegardnervirinae_like | 1.48 |
| Myoviridae | 1.43 |
| Schitoviridae | 0.91 |
| Arenaviridae_like | 0.63 |
| Autographiviridae | 0.54 |
| Queuovirinae | 0.52 |
| Herelleviridae | 0.48 |
| Podoviridae | 0.34 |
| Deejayvirinae | 0.27 |
| Gladiatorvirus | 0.21 |
| Demerecviridae | 0.19 |
| Casadabanvirus_like | 0.15 |
| Bronfenbrennervirinae | 0.06 |
| Guernseyvirinae | 0.06 |
| Fiersviridae | 0.03 |
| Steigviridae | 0.03 |
| **ARN virus Family** | |
| Yueviridae | 71.2219 |
| Narnaviridae | 23.4696 |
| Dicistroviridae | 4.8938 |
| Nodaviridae | 0.273 |
| Iflaviridae | 0.1109 |
| Tombusviridae | 0.0121 |
| Weivirus | 0.0121 |
| Picornaviridae | 0.0036 |
| Qinviridae | 0.001 |
| Spinareovirinae | 0.0004 |
| Zhaovirus | 0.0003 |
| Phenuiviridae | 0.0003 |
| Flaviviridae | 0.0003 |
| Yanvirus | 0.0003 |
| Hepeviridae | 0.0002 |
| Luteoviridae | 0.0002 |

**Table S1.** DNA and RNA virus family abundance (%).

| DNA virus Family | Abundance (%) |
|---|---|
| Peduoviridae | 54.58 |
| Family_unknown | 22.81 |
| Adenoviridae_like | 7.2 |
| Siphoviridae | 3.43 |
| Inoviridae_like | 2.81 |
| Zobellviridae | 1.82 |
| Gracegardnervirinae_like | 1.48 |
| Myoviridae | 1.43 |
| Schitoviridae | 0.91 |
| Arenaviridae_like | 0.63 |
| Autographiviridae | 0.54 |
| Queuovirinae | 0.52 |
| Herelleviridae | 0.48 |
| Podoviridae | 0.34 |
| Deejayvirinae | 0.27 |
| Gladiatorvirus | 0.21 |
| Demerecviridae | 0.19 |
| Casadabanvirus_like | 0.15 |
| Bronfenbrennervirinae | 0.06 |
| Guernseyvirinae | 0.06 |
| Fiersviridae | 0.03 |
| Steigviridae | 0.03 |
| **ARN virus Family** | |
| Yueviridae | 71.2219 |
| Narnaviridae | 23.4696 |
| Dicistroviridae | 4.8938 |
| Nodaviridae | 0.273 |
| Iflaviridae | 0.1109 |
| Tombusviridae | 0.0121 |
| Weivirus | 0.0121 |
| Picornaviridae | 0.0036 |
| Qinviridae | 0.001 |
| Spinareovirinae | 0.0004 |
| Zhaovirus | 0.0003 |
| Phenuiviridae | 0.0003 |
| Flaviviridae | 0.0003 |
| Yanvirus | 0.0003 |
| Hepeviridae | 0.0002 |
| Luteoviridae | 0.0002 |

**Table S3. RNA Virus Diversity.**

| Samples | Shannon | Simpson | Observed |
|---|---|---|---|
| Baseline_1 | 0.6931295 | 0.3573803 | 36 |
| Baseline_2 | 0.7817699 | 0.4173796 | 39 |
| Baseline_3 | 0.8051356 | 0.4261885 | 48 |
| Baseline_4 | 0.7813589 | 0.4360041 | 34 |
| Baseline_5 | 0.6586159 | 0.3114876 | 38 |
| Baseline_6 | 0.6534136 | 0.3326581 | 16 |
| Baseline_7 | 0.7541466 | 0.3924601 | 45 |
| Baseline_8 | 0.6479553 | 0.3191869 | 33 |
| HM_1 | 0.8468678 | 0.5078805 | 19 |
| HM_2 | 0.8458475 | 0.5200172 | 17 |
| HM_3 | 0.8347911 | 0.4923641 | 38 |
| HM_4 | 0.628322 | 0.316761 | 21 |
| HM_5 | 0.8227185 | 0.4901665 | 19 |
| HM_6 | 0.7875997 | 0.4651599 | 16 |
| HM_7 | 0.8531984 | 0.4876972 | 37 |
| HM_8 | 0.8200022 | 0.5264652 | 13 |
| HMM_1 | 0.7545611 | 0.423121 | 28 |
| HMM_2 | 0.5945537 | 0.2595464 | 50 |
| HMM_3 | 0.7792872 | 0.420593 | 40 |
| HMM_4 | 0.7286409 | 0.3595241 | 55 |
| HMM_5 | 0.7271032 | 0.3964098 | 17 |
| HMM_6 | 0.8793862 | 0.4772986 | 89 |
| HMM_7 | 0.8369004 | 0.4775304 | 46 |
| HMM_8 | 0.8321624 | 0.4657536 | 42 |
| LM_1 | 0.743583 | 0.3796968 | 47 |
| LM_2 | 0.849285 | 0.5125291 | 22 |
| LM_3 | 0.8217947 | 0.4699756 | 27 |
| LM_4 | 0.7368764 | 0.3867605 | 28 |
| LM_5 | 0.6428101 | 0.3035472 | 44 |
| LM_6 | 0.7029973 | 0.3601242 | 44 |
| LM_7 | 0.7553341 | 0.4037933 | 41 |

**Table S4. Regression model DNA viruses and environmental variables.**
glm.nb(formula = Virus ~ meiofauna + macrof + Bacteria + CH4 + OM + DIC_Flux + O2_Flux, data = combined_data, na.action = na.omit, init.theta = 98.03411864, link = log)

| | Estimate | Std. Error | z value | Pr(>\|z\|) | |
|---|---|---|---|---|---|
| **Intercept** | 4.99E+00 | 7.07E-01 | 7.062 | 1.64E-12 | *** |
| **meiofauna** | 1.37E-02 | 3.30E-03 | 4.146 | 3.38E-05 | *** |
| **macrofauna** | 1.30E+00 | 2.64E-01 | 4.917 | 8.81E-07 | *** |
| **Bacteria-activity** | 3.09E-08 | 1.44E-08 | 2.147 | 0.03178 | * |
| **$CH_4$** | -1.40E-02 | 4.63E-03 | -3.013 | 0.00258 | ** |
| **OM%** | 9.95E-02 | 4.36E-02 | 2.281 | 0.02256 | * |
| **DIC_Flux** | 1.17E-04 | 6.67E-05 | 1.752 | 0.0798 | . |
| **$O_2$_Flux** | -1.11E-03 | 1.59E-04 | -6.978 | 2.99E-12 | *** |

Signif. codes: 0 ‘***’ 0.001 ‘**’ 0.01 ‘*’ 0.05 ‘.’ 0.1 ‘ ’ 1

**Table S5. DNA Viruses.** McFadden's $R^2$ values.

| | P-values | McFadden’s $R^2$ |
|---|---|---|
| **Intercept** | 1.64E-12 | 0.7930 |
| **Meiofauna** | 3.38E-05 | 0.1257 |
| **Bacteria-activity** | 3.18E-02 | 0.0064 |
| **Macrofauna** | 8.81E-07 | 0.0001 |
| **$CH_4$** | 2.58E-03 | 0.0635 |
| **OM%** | 0.02256219 | 0.0003 |
| **DIC_Flux** | 7.98E-02 | 0.0023 |
| **$O_2$_Flux** | 2.99E-12 | 0.0892 |

**Table S6. DNA viruses and variables. Adonis (PERMNOVA) analysis.**

| | Df | SumOfSqs | R2 | F | Pr(>F) |
|---|---|---|---|---|---|
| **Meiofauna** | 1.00E+00 | 5.18E-02 | 0.17742535 | 6.72E+00 | 0.015 |
| **Macrofauna** | 1.00E+00 | 6.32E-02 | 0.21639537 | 8.192657 | 0.003 |
| **Bacteria-activity** | 1.00E+00 | 1.41E-02 | 0.04815896 | 1.823282 | 0.202 |
| **$CH_4$** | 1.00E+00 | 2.58E-02 | 0.08832139 | 3.34E+00 | 0.074 |
| **OM%** | 1.00E+00 | 1.48E-02 | 0.05079208 | 1.922971 | 0.176 |
| **DIC_Flux** | 1.00E+00 | 1.03E-02 | 0.03546133 | 1.342554 | 0.268 |
| **$O_2$_Flux** | 1 | 0.12883066 | 0.44144819 | 16.713083 | 0.001 |
| **Residual** | 10 | 0.07708372 | 0.26413331 | NA | NA |
| **Total** | 17 | 0.29183642 | 1 | NA | NA |

**Table S7. Regression model RNA viruses and environmental variables.**
glm.nb(formula = Virus ~ meiofauna + macrof + Bacteria + CH4 + OM + DIC_Flux + O2_Flux, data = combined_data, na.action = na.omit, init.theta = 98.03411864, link = log)

| | Estimate | Std. Error | z value | Pr(>\|z\|) | |
|---|---|---|---|---|---|
| **Intercept** | 1.02E+01 | 2.79E-01 | 36.454 | 2.00E-16 | *** |
| **meiofauna** | -2.96E-04 | 1.45E-03 | -0.204 | 8.38E-01 | |
| **macrofauna** | -9.37E-03 | 9.63E-02 | -0.097 | 9.23E-01 | |
| **Bacteria-activity** | 2.94E-08 | 5.91E-09 | 4.968 | 6.78E-07 | *** |
| **$CH_4$** | 3.93E-03 | 2.03E-03 | 1.936 | 0.05291 | |
| **OM%** | 3.50E-02 | 1.75E-02 | 1.999 | 0.0456 | * |
| **DIC_Flux** | 9.78E-05 | 2.96E-05 | 3.302 | 0.00096 | *** |
| **$O_2$_Flux** | -6.58E-05 | 5.62E-05 | -1.171 | 2.41E-01 | |

**Table S8. RNA viruses McFadden's $R^2$ values**

| | P-values | McFadden's $R^2$ |
|---|---|---|
| **Intercept** | 5.98E-291 | 8.09E-01 |
| **meiofauna** | 8.38E-01 | 0.0489 |
| **Bacteria-activity** | 6.78E-07 | 0.5698 |
| **macrofauna** | 9.23E-01 | 0.0496 |
| **$CH_4$** | 5.29E-02 | 0.0064 |
| **OM%** | 4.56E-02 | 0.0655 |
| **DIC_Flux** | 9.60E-04 | 0.1330 |
| **$O_2$_Flux** | 2.41E-01 | 0.0305 |

**Table S9. RNA viruses and variables. Adonis (PERMNOVA) analysis.**

| | Df | SumOfSqs | R2 | F | Pr(>F) |
|---|---|---|---|---|---|
| **Meiofauna** | 1 | 0.000734 | 0.004734 | 0.122271 | 0.87 |
| **Macrofauna** | 1 | 0.000561 | 0.003618 | 0.093451 | 0.878 |
| **Bacteria-activity** | 1 | 0.029881 | 0.192749 | 4.978281 | 0.027 |
| **$CH_4$** | 1 | 0.031218 | 0.201370 | 5.200927 | 0.032 |
| **OM%** | 1 | 0.003234 | 0.020860 | 0.538756 | 0.579 |
| **DIC_Flux** | 1 | 0.027032 | 0.174372 | 4.503642 | 0.046 |
| **$O_2$_Flux** | 1 | 0.003725 | 0.024025 | 0.620523 | 0.535 |

**Table S10. Meiofauna abundance.**

| Samples | Nematoda | Ostracoda | Copepoda | Kinorhyncha | Turbellaria | Halacaridae | Oligochaeta | Bosmina | Monoporeia | Platyhelminthes |
|---|---|---|---|---|---|---|---|---|---|---|
| Baseline_1 | 74 | 9 | 86 | 0 | 5 | 0 | 0 | 0 | 0 | 0 |
| Baseline_2 | 29 | 1 | 33 | 1 | 2 | 2 | 0 | 0 | 0 | 0 |
| Baseline_3 | 115 | 56 | 49 | 0 | 4 | 0 | 0 | 3 | 0 | 0 |
| Baseline_4 | 58 | 2 | 2 | 0 | 1 | 0 | 0 | 4 | 0 | 1 |
| Baseline_5 | 55 | 5 | 104 | 0 | 2 | 1 | 0 | 0 | 2 | 0 |
| Baseline_7 | 70 | 15 | 43 | 1 | 4 | 0 | 0 | 1 | 1 | 0 |
| Baseline_8 | 52 | 10 | 41 | 0 | 3 | 1 | 0 | 0 | 0 | 0 |
| HM 1 | 13 | 15 | 10 | 1 | 0 | 0 | 0 | 0 | 0 | 0 |
| HM 2 | 135 | 85 | 5 | 0 | 3 | 0 | 0 | 0 | 0 | 0 |
| HM 3 | 80 | 32 | 10 | 0 | 0 | 1 | 0 | 0 | 0 | 0 |
| HM 4 | 84 | 3 | 24 | 2 | 0 | 1 | 0 | 1 | 0 | 0 |
| HM 5 | 42 | 22 | 4 | 1 | 0 | 0 | 0 | 0 | 0 | 0 |
| HM 6 | 40 | 6 | 11 | 0 | 1 | 1 | 0 | 5 | 0 | 0 |
| HM 7 | 22 | 1 | 3 | 0 | 0 | 0 | 0 | 0 | 0 | 0 |
| HMM 1 | 51 | 1 | 17 | 0 | 0 | 0 | 0 | 4 | 0 | 0 |
| HMM 2 | 55 | 14 | 3 | 0 | 1 | 0 | 0 | 0 | 0 | 0 |
| HMM 3 | 30 | 8 | 16 | 1 | 1 | 1 | 0 | 0 | 0 | 0 |
| HMM 5 | 59 | 5 | 19 | 1 | 1 | 0 | 0 | 6 | 0 | 0 |
| HMM 6 | 23 | 1 | 6 | 0 | 0 | 0 | 0 | 1 | 0 | 0 |
| HMM 7 | 156 | 3 | 24 | 1 | 0 | 2 | 0 | 16 | 0 | 0 |
| HMM 8 | 29 | 9 | 14 | 0 | 0 | 1 | 0 | 0 | 1 | 0 |
| LM 1 | 7 | 2 | 0 | 0 | 0 | 0 | 0 | 0 | 0 | 0 |
| LM 2 | 13 | 11 | 4 | 0 | 0 | 0 | 0 | 0 | 0 | 0 |
| LM 3 | 10 | 6 | 5 | 0 | 0 | 0 | 0 | 0 | 0 | 0 |
| LM 5 | 37 | 11 | 7 | 0 | 0 | 0 | 0 | 0 | 0 | 0 |
| LM 6 | 17 | 5 | 14 | 0 | 1 | 0 | 0 | 0 | 0 | 0 |
| LM 7 | 11 | 0 | 11 | 0 | 0 | 0 | 0 | 1 | 1 | 0 |
| LM 8 | 4 | 3 | 6 | 0 | 0 | 0 | 0 | 0 | 0 | 0 |

Meiofauna were normalize to volume samples

| Samples | Sediment volume (ml) |
|---|---|
| Baseline_1 | 4 |
| Baseline_2 | 3 |
| Baseline_3 | 8 |
| Baseline_4 | 5 |
| Baseline_5 | 5 |
| Baseline_7 | 7 |
| Baseline_8 | 5 |
| HM 1 | 3 |
| HM 2 | 8 |
| HM 3 | 7 |
| HM 4 | 5 |
| HM 5 | 7 |
| HM 6 | 4 |
| HM 7 | 2 |
| HMM 1 | 5 |
| HMM 2 | 3 |

| | |
|---|---|
| HMM 3 | 3 |
| HMM 5 | 3 |
| HMM 6 | 3 |
| HMM 7 | 5 |
| HMM 8 | 5 |
| LM 1 | 4 |
| LM 2 | 4 |
| LM 3 | 4 |
| LM 5 | 10 |
| LM 6 | 8 |
| LM 7 | 8 |
| LM 8 | 3 |

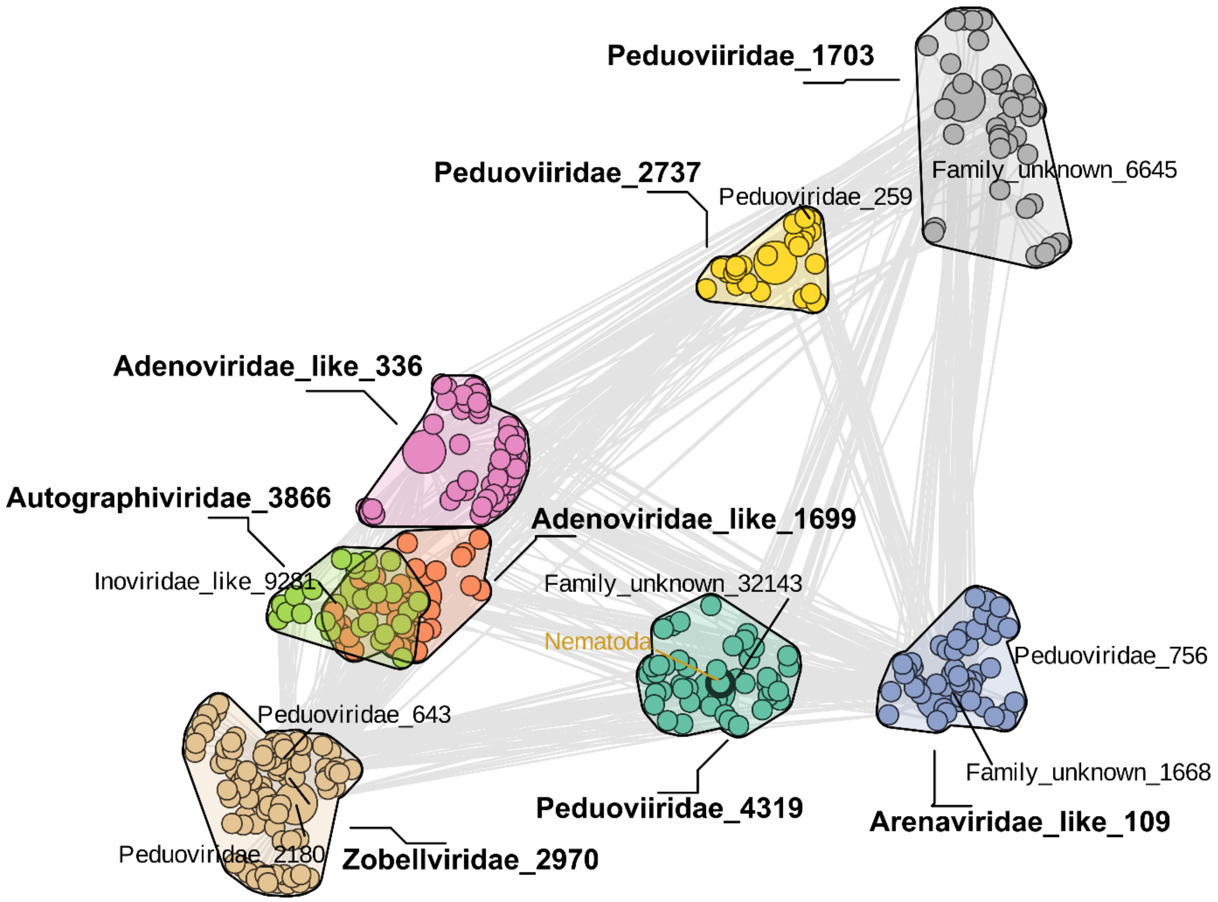

**Fig. S1. Co-abundance of DNA viruses, bacteria (activity) and meiofauna.** Nematoda appeared as significant.

Fig.

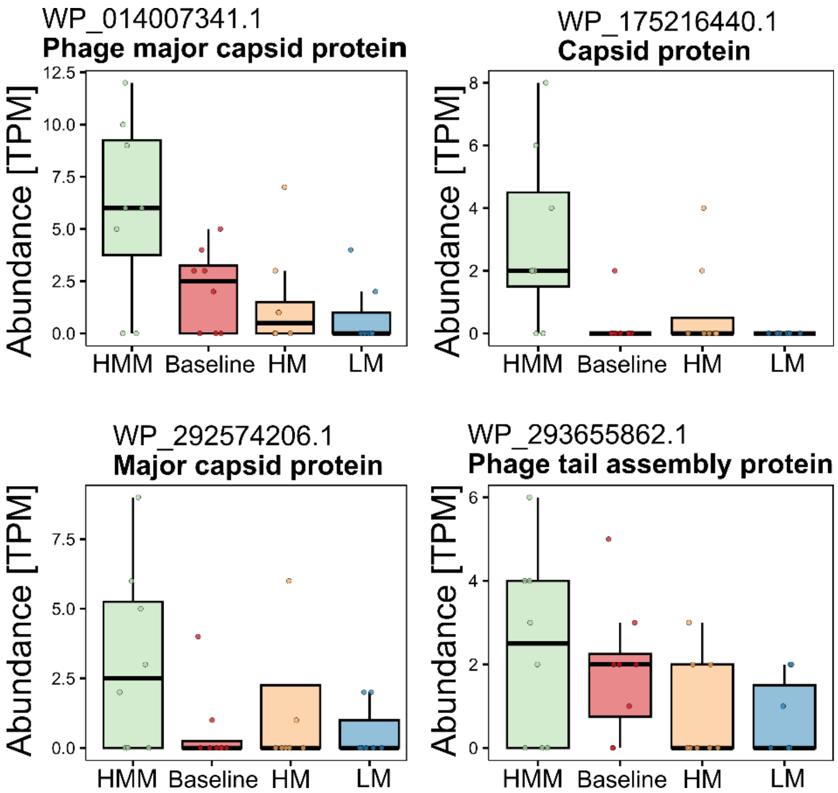

**S2. Gene expression of four contrasting viral genes (capsid and tail) in infauna treatments.**

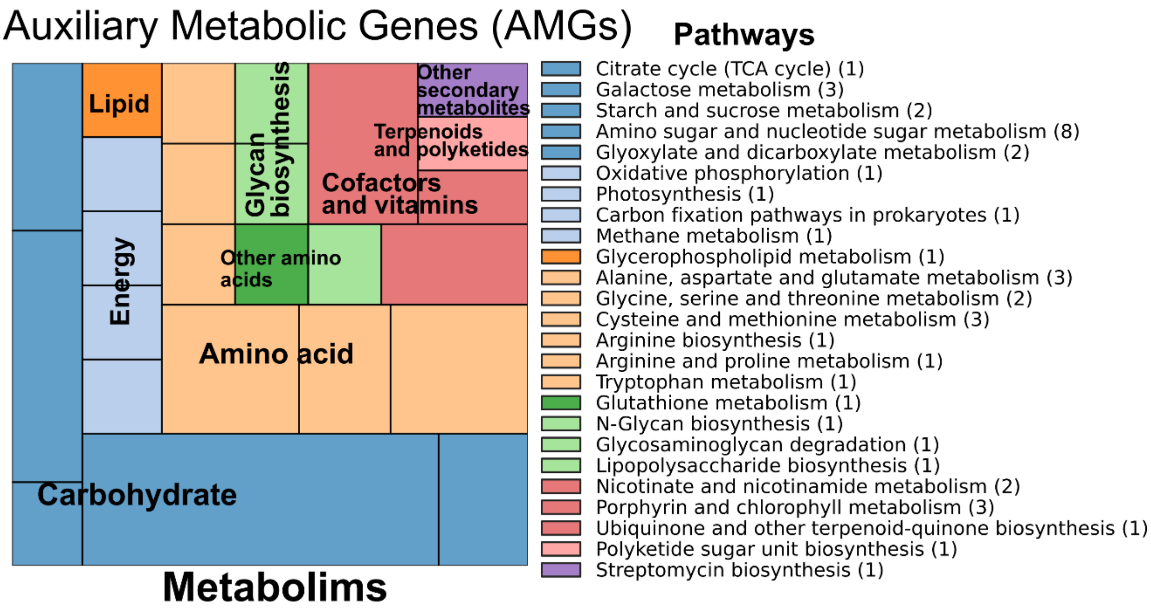

**Fig. S3. Auxiliary metabolic genes found in the genomes of the DNA virus by pathway and metabolism.**
